# Advances in non-invasive tracking of wave-type electric fish in natural and laboratory settings

**DOI:** 10.1101/2022.06.02.494479

**Authors:** Till Raab, Manu S. Madhav, Ravikrishnan P. Jayakumar, Jörg Henninger, Noah J. Cowan, Jan Benda

## Abstract

Recent technological advances greatly improved the possibility to study freely behaving animals in natural conditions. However, many systems still rely on animal-mounted devices, which can already bias behavioral observations. Alternatively, animal behaviors can be detected and tracked in recordings of stationary sensors, e.g. video cameras. While these approaches circumvent the influence of animal-mounted devices, identification of individuals is much more challenging. We take advantage of the individual-specific electric fields electric fish generate by discharging their electric organ (EOD) to record and track their movement and communication behaviors without interfering with the animals themselves. EODs of complete groups of fish can be recorded with electrode arrays submerged in the water and then be tracked for individual fish. Here, we present an improved algorithm for tracking electric signals of wave-type electric fish with unprecedented accuracy. Our algorithm benefits from combining and refining previous approaches of tracking individual specific EOD frequencies (EODf) and spatial electric field properties. In this process, the similarity of signal pairs in extended data windows determines their tracking order, making the algorithm more robust against detection losses and intersections. We quantify the performance of the algorithm and show its application for a data set recorded with a 64-electrode array in a stream in the Llanos, Colombia, where we managed, for the first time, to track *Apteronotus leptorhynchus* over many days. These technological advances make electric fish a unique model system for a detailed analysis of social and communication behaviors, with strong implications for our research on sensory coding.

## 1 INTRODUCTION

Unraveling causal factors driving various animal behaviors in experimental and in particular in observational studies is challenging, since most behaviors result from an integration of a broad range of social and environmental stimuli, internal states, and past experiences (Chapman et al., 1995; Sapolsky, 2005; Boon et al., 2007; Markham et al., 2015). In laboratory studies, environments and contexts are systematically simplified in order to minimize the number of potential factors influencing behaviors (e.g. Pantoni et al., 2020; Bastian et al., 2001). Such studies are tailored to specific behaviors and well defined contexts. However, behaviors in such constrained settings often deviate from behaviors in natural environments and thus have to be interpreted with care (Cheney et al., 1995; Rendall et al., 1999; Henninger et al., 2018). To discover behavioral traits of interest in the first place, field studies or laboratory experiments with complex, more naturalistic designs are needed. Recent technological advances in remote recording techniques, tags, and data loggers, as well as advances in data analysis, facilitate the collection and evaluation of comprehensive and viable data in naturalistic settings with freely moving and interacting animals (Dell et al., 2014; Hughey et al., 2018; Mathis et al., 2018; Jolles, 2021). These new big-data approaches open up new opportunities in behavioral research in that they potentially allow to quantitatively study animal behaviors in more complex and naturalistic settings (Gomez-Marin et al., 2014; Egnor and Branson, 2016).

A suitable recording technique can be selected from a large variety of available devices to match the requirements imposed by the model species, environmental conditions, and the scientific question (Hughey et al., 2018). Animal mounted bio-loggers are a commonly used technique (Nagy et al., 2010; Strandburg-Peshkin et al., 2017; Hughey et al., 2018). Different types of sensors allow for studying various aspects of animal behaviors across species (e.g. Nagy et al., 2010; Robinson et al., 2012; Strandburg-Peshkin et al., 2015, 2018. However, bio-loggers require frequent animal handling and animals are required to carry devices, both inducing a potential bias (Saraux et al., 2011). Furthermore, bio-loggers might miss relevant information, since not all interacting animals might be equipped with a logger (e.g. Strandburg-Peshkin et al., 2019), signal detection range is limited, or data are recorded discontinuously to extend the overall recording period (Strandburg-Peshkin et al., 2017; Hughey et al., 2018).

Alternatively, behaving animals can be tracked by means of remote sensing devices (Kühl and Burghardt, 2013; Theriault et al., 2014; Henninger et al., 2018; Hughey et al., 2018; Torney et al., 2018; Raab et al., 2019; Henninger et al., 2020; Aspillaga et al., 2021). In this approach, recorded signals can origin from small micro-transmitters that get affixed to animals (e.g. acoustic telemetry system for fish: Aspillaga et al., 2021) or from the animals themselves (photography, video recordings: Sherley et al., 2010; Lahiri et al., 2011; Theriault et al., 2014; Nourizonoz et al., 2020, ultrasound vocalizations: Surlykke and Kalko, 2008; Seibert et al., 2013; Hügel et al., 2017, electric signals: Henninger et al., 2018; Raab et al., 2019; Fortune et al., 2020). These methods benefit from minimal interference with the animals themselves. On the other hand, covering large areas is costly. In particular, tracking animal identity can be quite challenging (“biometrics”, Kühl and Burghardt, 2013) and requires sophisticated and computationally demanding pre-processing of the data. If individuals do not have specific invariant characteristics, like, for example, the stripes of a zebra (Lahiri et al., 2011), then tracking algorithms need to handle temporally changing biometric profiles that often overlap in their characteristics (e.g. Madhav et al., 2018).

Electric fish are particularly well suited for being tracked in the laboratory and in their natural habitats based on remote sensing (Jun et al., 2013; Henninger et al., 2018; Madhav et al., 2018; Raab et al., 2019; Fortune et al., 2020). Wave-type electric fish produce a semi-sinusoidal electric field through continuous discharges of an electric organ (EOD, Turner et al., 2007) that they use for electrolocation (Fotowat et al., 2013) and communication (Albert and Crampton, 2005; Smith, 2013; Benda, 2020). The frequency of the EOD (EODf) is individual specific and remarkably stable over minutes to hours (Moortgat et al., 1998), providing a characteristic biometric cue. The EODs of many electric fish can be recorded by means of an array of submerged electrodes without the need to catch and tag the fish (Henninger et al., 2018). From these recordings, individual fish have to be identified and tracked over time. Previous tracking approaches were either based on EODf (Henninger et al., 2020) or on spatial electric field properties that can be reconstructed from signal powers across recording electrodes (Madhav et al., 2018). However, the latter depends on the fish’s spatial position and orientation and EODf is sensitive to temperature changes (Dunlap et al., 2000) and is actively modulated for electrocommunication (Smith, 2013). Both tracking features might fail when fish are close by, either in their EODf or spatially, especially in recordings of electric fish in high densities.

In the following, we describe and evaluate an improved tracking algorithm for wave-type electric fish recorded with electrode arrays. By combining, refining, and extending previous approaches, our algorithm is capable of tracking EODs of individual fish with unprecedented accuracy. Since both movement behaviors (Madhav et al., 2018; Henninger et al., 2020) and communication (Smith, 2013; Henninger et al., 2018; Fortune et al., 2020) can be analyzed based on EOD recordings, our algorithm is a fundamental advancement for a wide range of behavioral studies on freely moving and interacting electric fish (Raab et al., 2019, 2021). Finally, we demonstrate the performance of our tracking algorithm on recordings of *Apteronotus leptorhynchus* taken with an array of 64 electrodes in a stream in the Llanos in Colombia.

## 2 MATERIALS AND EQUIPMENT

### 2.1 Data acquisition

EODs of freely swimming fish were recorded with arrays of monopolar electrodes at low-noise buffer headstages (1×gain, 10×5×5mm^3^, fig. 1B) arranged in grid-like structures (Fig. 1C, D). Electric signals are amplified (100×gain, 100Hz high-pass filter, 10 kHz low-pass), digitized at 20 kHz with 16 bit resolution, and stored on external data storage devices for later offline analysis. The custom-built recording systems (npi-electronics GmbH, Tamm, Germany) were powered by car batteries (12 V, 80 Ah). Various configurations of the electrode arrays have been successfully used to record populations of electric fish in the wild (Henninger et al., 2018, 2020, unpublished field-trips: Colombia 2016, 2019, fig. 1C, E), as well as in the laboratory (Raab et al., 2019, 2021, fig. 1D, F). The first 64-channel amplifier system with external computer for data storage (Henninger et al., 2018, 2020, Colombia 2016), data acquisition via two data acquisition boards (PCI-6259, National Instruments, Austin, Texas, USA) was controled by C++ software (https://github.com/bendalab/fishgrid). For the 2019 recordings in Colombia we used a modular 16-channel system based on a Raspberry Pi 3B (Raspberry Pi Foundation, UK) that stores the data digitized by an USB data acquisition board (USB-1608GX, Measurement Computing, Norton, MA, USA) on an 256 GB USB stick controlled by python software https://whale.am28.uni-tuebingen.de/git/raab/Rasp_grid.git (fig. 1A).

**Figure 1.**
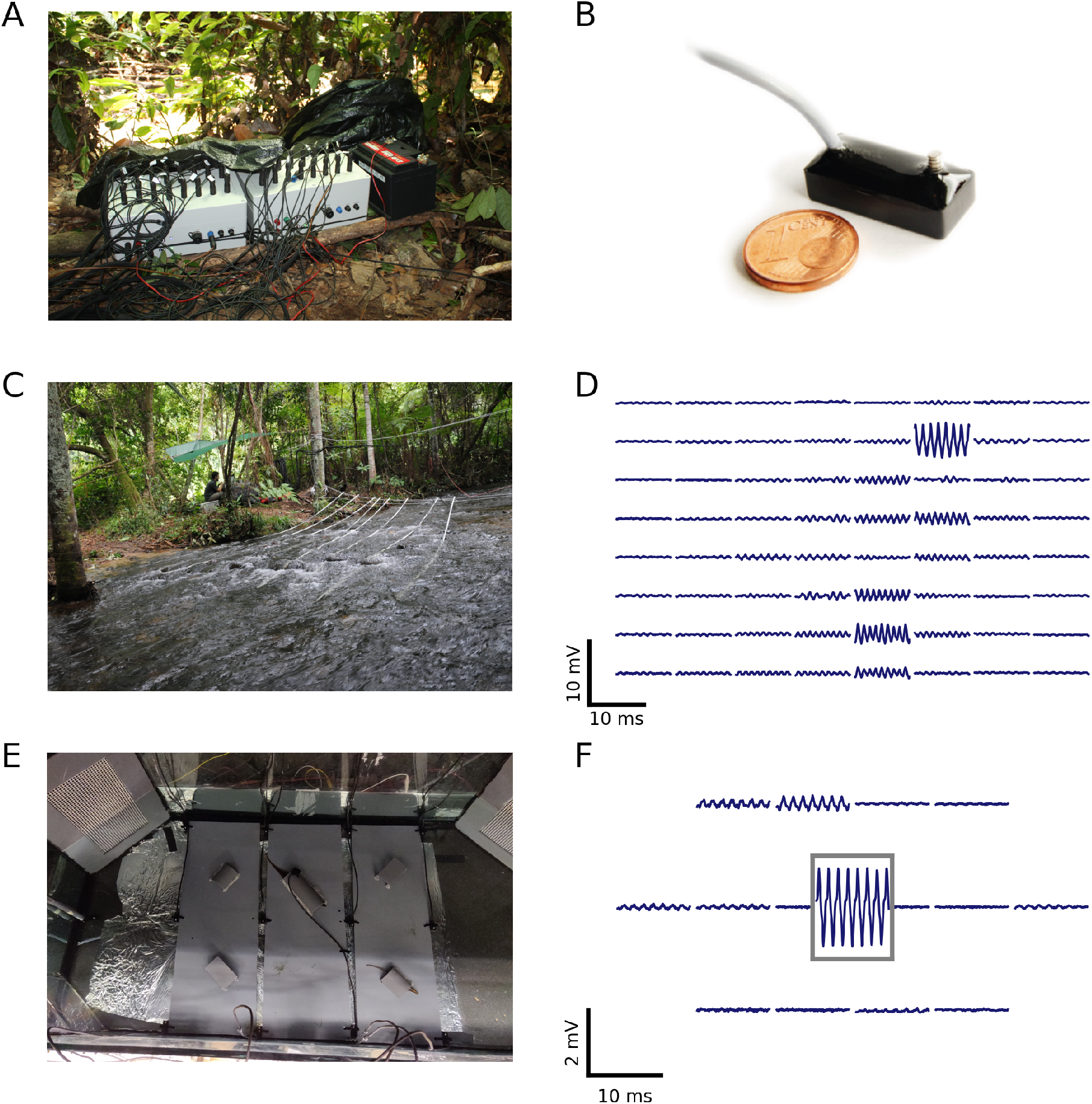
Recording systems, electrode arrangements, and corresponding signals of recorded electric fish. **A** Two of the Raspberry-based 16-channel amplifiers and recorders used for an array with 32 electrodes. **B** Monopolar stainless-steel electrode on headstage used for recordings in the field and laboratory experiments (after Henninger, 2015). **C** Recording setup used to record a population of *A. leptorhynchus* in the Rio Rubiano, Colombia, in 2016. 64 electrodes were mounted on PVC-tubes and arranged in an 8 × 8 grid covering an area of 3.5 × 3.5 m^2^. **D** Snapshot of the electric signals recorded with the setup shown in panel C. The top left panel corresponds to the most upstream electrode mounted on the tube closest to the river bank. **E** Recording setup used to record electric signals of pairs of *A. leptorhynchus* during competitions in a laboratory experiment (Raab et al., 2021). 15 electrodes were uniformly distributed at the bottom of the aquarium and one electrode was placed in the central tube the fish compete for. **F** Snapshot of electric signals recorded during the competition experiment shown in panel E. The signal framed in grey is from the central electrode located in the optimal tube. The EOD waveform shows the characteristic shoulder of *A. leptorhynchus*.

### 2.2 Spectrograms

EODs of individual fish are identified and extracted from the electric recordings based on their EODf and respective harmonic structure (Fig. 2C). For each electrode we compute power spectral densities (PSDs) of overlapping data snippets shifted by Δ*t* ≈ 300 ms (Fig. 2 A). The size of fast Fourier transform (FFT) windows was set to *n*_fft_ = 2^15^ ≈ 1.64 s (e.g. Raab et al., 2021) or *n*_fft_ = 2^16^ ≈ 3.28 s (e.g. Raab et al., 2019; field recordings displayed in fig. 9) to result in frequency resolutions of 0.6 Hz and 0.3 Hz, respectively, needed to resolve EODfs in high fish densities.

**Figure 2.**
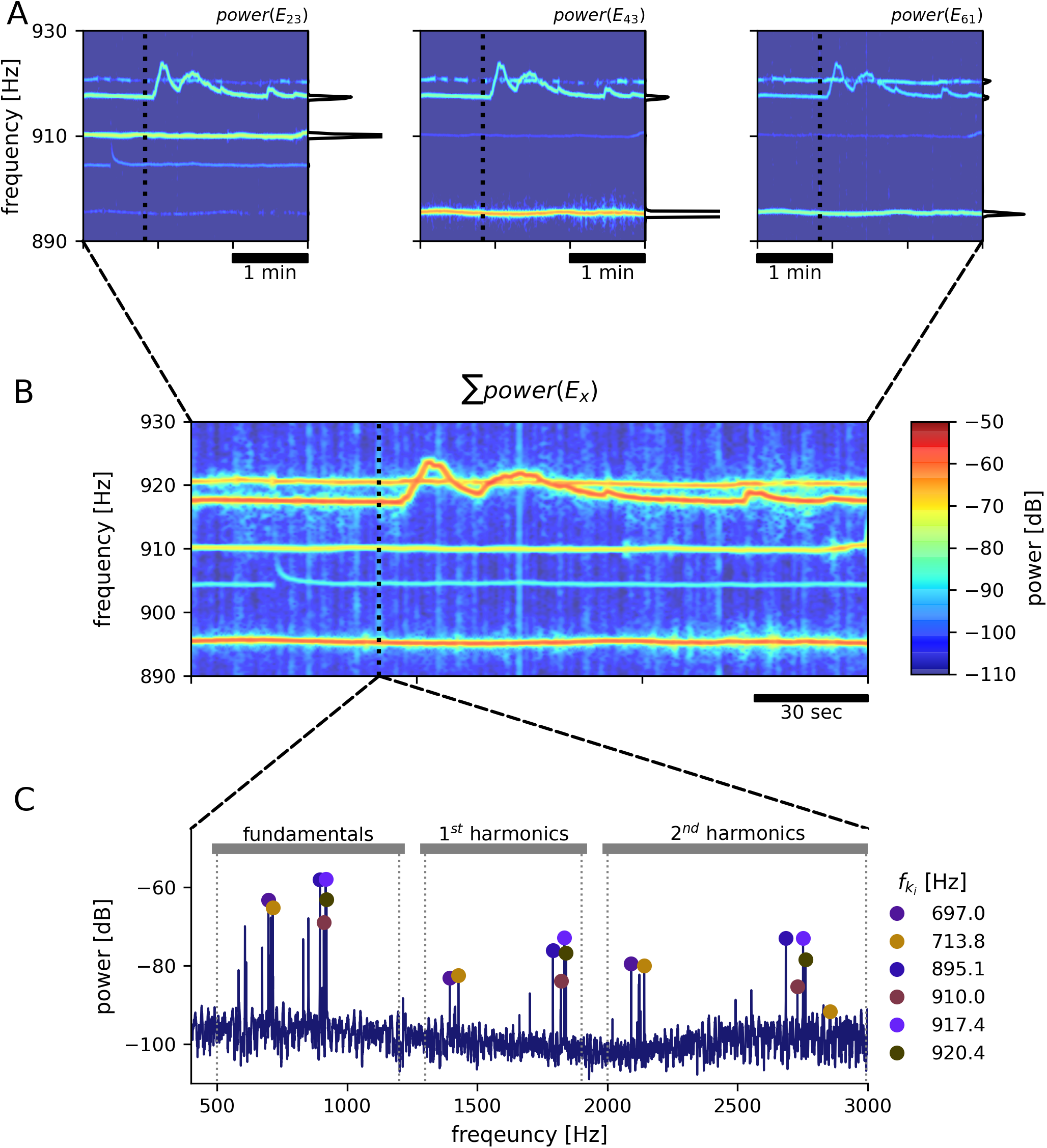
EOD frequency extraction from recordings with an electrode array. As an example, a 3 min snippet of a recording with the 8 × 8 array from Rio Rubiano, Colombia, taken during the day of April 10^th^, 2016 is shown. **A** Spectrograms from three different electrodes. Warmer colors represent increased power in respective frequencies. EODfs of individual *A. leptorhynchus* remain rather stable, except during electrocommunication (e.g. EODf trace starting at ~ 917 Hz). A non-logarithmic PSD extracted at time 50 s indicated by the dotted line is shown at the side of each panel. **B** The summed up spectrogram over all electrodes contains distinct traces from many different fish. **C** Peaks are detected in the summed up power spectra that are then clustered into frequency groups of a fundamental frequency and at least two of its harmonics, corresponding to a specific fish (Henninger et al., 2020). Fundamental EOD frequenciess, their corresponding powers in each electrode and their detection times are stored for subsequent tracking.

### 2.3 Extraction of EODf and feature vector

In order to detect EODfs of all recorded fish, for each time point *t_i_* PSDs from all electrodes were summed up (Fig. 2 B). The summed PSDs were transformed to decibel levels, *L*(*f*) = 10 log_10_(*P*(*f*)/*P*_0_), relative to a power of *P*_0_ = 1 mV^2^/Hz. In these logartihmic power spectra, peaks were detected (Todd and Andrews, 1999) and groups of harmonics were assigned to their corresponding fundamental frequencies (fig. 2C). See Henninger et al. (2020) and the harmonics.py module in the thunderfish package (https://github.com/bendalab/thunderfish) for details.

Harmonic groups were extracted from the summed power spectra in order to save computing time. Extracting fundamental frequencies from each of *n* electrodes separately would take n-times longer, but might be more advantageous for separating distant fish that are close by in EODf. We are therefore working on improving the performance of the harmonic-group extraction. The tracking algorithm described in the Method section is independent of whether fundamental frequencies were obtained from the individual spectra or the summed one.

For each time point *t_i_* and each signal indexed by *k*, a feature vector

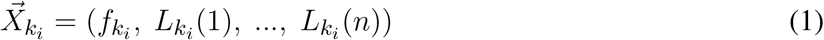

is assembled that includes the fundamental EODf, *f_k_i__*, and the corresponding logarithmic powers, *L_k_i__* (*x*), in the PSDs of all *n* recording electrodes *x*. Based on this feature vector the individual fish are tracked as described in the following Methods.

## 3 METHODS

In the following we present an algorithm for tracking wave-type electric fish in electrode-array recordings. The algorithm merges and extends two complementary approaches that are based on EODf (Henninger et al., 2018, 2020) or on the spatial distribution of signal powers (Madhav et al., 2018). We then test the performance of the tracking algorithm against manually tracked data. Open-source Python scripts for tracking and post-processing of analyzed data can be obtained from https://github.com/bendalab/wavetracker.

### 3.1 Algorithm for tracking wave-type electric fish

Both EODf and the spatial distribution of EOD power across electrodes change with time and potentially overlap between fish. EODf can be actively altered in the context of communication (Smith, 2013; Benda, 2020) and the signal powers across electrodes change with the fish’s motion (Madhav et al., 2018). This variability and potential overlap in signal features challenges reliable tracking, especially in recordings with many fish.

Furthermore, the existing algorithms track signals in the order of their temporal detection, i.e. signals detected in consecutive time steps are directly assigned to already tracked EODf traces (Madhav et al., 2018; Henninger et al., 2020). Potentially this leads to tracking errors, because even with the utilization of an electrode array, EODs of freely moving and interacting electric fish are rarely detected continuously, i.e. consecutively in subsequent time steps. Low signal-to-noise ratios, resulting from large distances between fish and recording electrodes or objects like rocks or logs distorting or even blocking electric fields, frequently lead to detection losses. When multiple fish with similar EODfs are recorded simultaneously, EODf traces can potentially cross each other (e.g. in the context of emitted communication signals, Benda, 2020). Especially in these occasions, detection losses can frequently result in tracking errors.

In order to improve on these issues, we developed a tracking algorithm which, first, is based on feature vectors that include both EODf and signal power across electrodes (Fig. 3) and, second, is less constrained by the temporal sequence of detected signals (Fig. 5).

**Figure 3.**
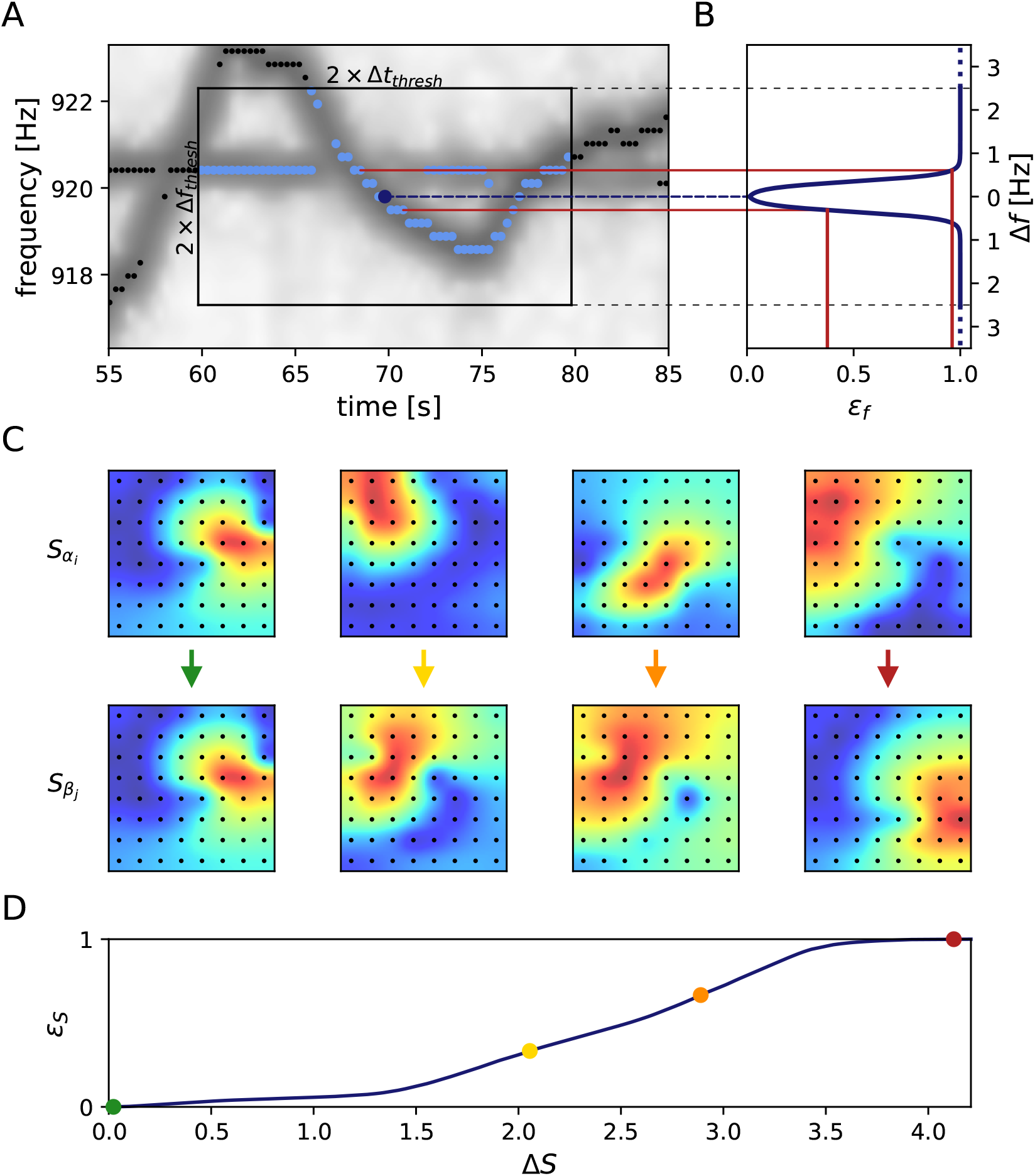
Frequency and field errors. **A** Summed spectrogram of a 30 s long part of the recording shown in Fig. 2B. For each electric fish signal, potential connection partners are limited by a time difference threshold, Δ*t_thresh_* = 10 s, and a frequency difference threshold, Δ*f_thresh_* = 2.5 Hz. For a given signal *α* with EODf *f_α_i__*. at time step *i* (dark blue dot), potential connection candidates *β* at different times *j* (light blue dots) need to be within these thresholds (box), whereas signals beyond these thresholds (black dots) are not considered. **B** Absolute frequency differences, Δ*f* Eq. (2), are mapped (red lines) to frequency errors, *ε_f_*, using a logistic function, Eq. (4) (line), favoring small frequency differences. **C** The field error as the second tracking parameter in addition to the frequency error is based on spatial profiles, Eq. (5), of signal powers over all electrodes (black dots). The field difference, Δ*S*, is computed as the Euclidean distance, Eq. (6), between the spatial profiles, Eq. (5), of potential signal pairs (columns). With decreasing similarity (columns left to right) the field difference increases. **D** To obtain normalized field errors, *ε_S_*, in a range similar to the one of the frequency errors, *ε_f_*, each field difference is set into perspective to a representative cumulative distribution (Eq. (7), black line) of field differences obtained by collecting all potential field differences of a manually selected 30 s window in the recording. The examples from panel C are marked by respective dots.

#### 3.1.1 Distance measure

We start out with extracting feature vectors 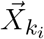, Eq. (1), containing an EOD frequency, *f_k_i__*, and its powers, *L_k_i__* (*x*), on all electrodes *x*, for all signals *k* and each time step *i*. In a first step the distance between all pairs of feature vectors, 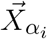 and 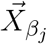, of signals *α* and *β* at times *i* ≠ *j* are quantified. Only pairs within a time difference of |*t_j_* – *t_i_*| ≤ Δ*t_thresh_* = 10 s and a maximum difference

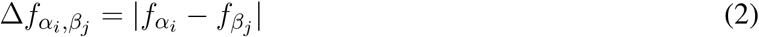

of Δ*f_thresh_* = 2.5 Hz between the two EOD frequencies of the feature vectors are considered (Fig. 3A).

The distance between the two signals *α_i_* and *β_j_*

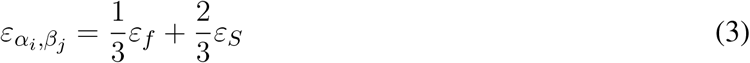

is computed as a weigthed sum of the frequency error, *ε_f_*, and the field error, *ε_S_*. Both errors range from 0 to 1 and are explained in the following sections. The field error gets twice the weigth of the frequency error, because tracking issues usually arise from low frequency errors. Nevertheless, the frequency error remains a relevant tracking feature, especially when fish are in close proximity resulting in low field errors.

#### 3.1.2 Frequency error

The frequency error is based on the difference in EOD frequencies, Eq. (2) and has been used previously to track signals of electric fish (Henninger et al., 2018, 2020). We transform the EODf difference, Eq. (2), into the frequency error

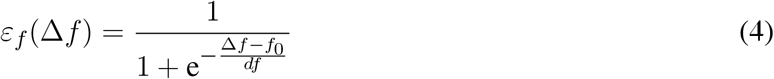

via a logistic function, that maps the EODf difference, Δ*f*, onto the interval from zero to one. The turning point of the logistic function at *f*_0_ = 0.35 Hz and the corresponding inverse slope, *df* = 0.08 Hz ensure a maximum frequency error already at small EODf differences of no more than about 0.8 Hz (Fig. 3 B). This transformation mitigates very small frequency differences and equalizes larger frequency differences in the assessment of whether two signals *α* and *β* originate from the same or different fish.

#### 3.1.3 Field error

EODf traces of electric fish occasionally cross each other, e.g. when individuals actively alter their EODf in the context of communication (e.g. Zupanc, 2002; Triefenbach and Zakon, 2008; Raab et al., 2021, fig. 3A). In these situations, frequency as a tracking feature fails. This is where the spatial properties of a signal, i.e. signal powers across recording electrodes that reflect the position and orientation of a fish, come into play (Madhav et al., 2018, fig. 3C). The signal powers, *L_k_i__* (*x*), are rescaled to the spatial profile

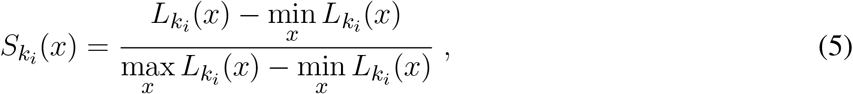

ranging between 0 and 1, for the smallest and largest power of that signal, respectively.

The field difference Δ*S*, i.e. the difference between the spatial profiles of two signals *α* and *β* at times *i* and *j*, is computed as their Euclidean distance according to

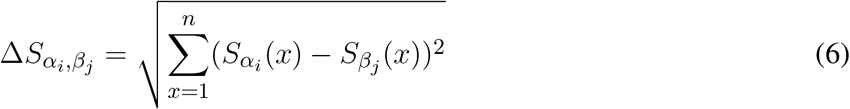

However, the magnitude of this difference depends on the configuration of the electrode array, especially on the number of recording electrodes. To obtain field errors, *ε_S_*, that are independent of electrode configuration, we map the field differences through their cumulative distribution:

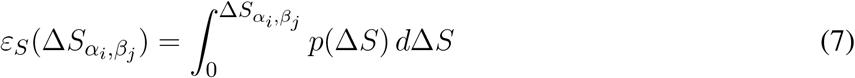

A representative distribution of field differences, *p*(Δ*S*), is estimated from the field differences between each potential signal pair (Δ*t_i,j_* ≤ ± 10 s, any frequency difference) within a 30 second data snippet where fish can be assumed to be active, i.e. during night time (Fig. 3D).

#### 3.1.4 Tracking within a data window

Now we know how to quantify the distance *ε*, Eq. (3), between to signals and we can proceed with the actual tracking algorithm. Based on the distances, the algorithm decides which signal pairs belong together in order to track individual fish throughout a recording. Computing the distances between all pairs of signals of a recording at once, however, is not feasible. Instead we break down the tracking into tracking windows of 30 seconds at a time (Fig. 5). Within these tracking windows, we first compute the distances, *ε*, between each potential signal pair *α_i_* and *β_j_* and store them in a three-dimensional distance cube, where the first two dimensions refer to signals *α_i_* and *β_j_* and the third dimension to the time steps *i* where signals *α_i_* have been detected (Fig. 4). Note that in each time step *i* we have a different number of signals *α_i_* and, consequently, the number of elements in the second dimension, referring to signals *β_j_* from all time steps *j* > *i* is also variable.

**Figure 4.**
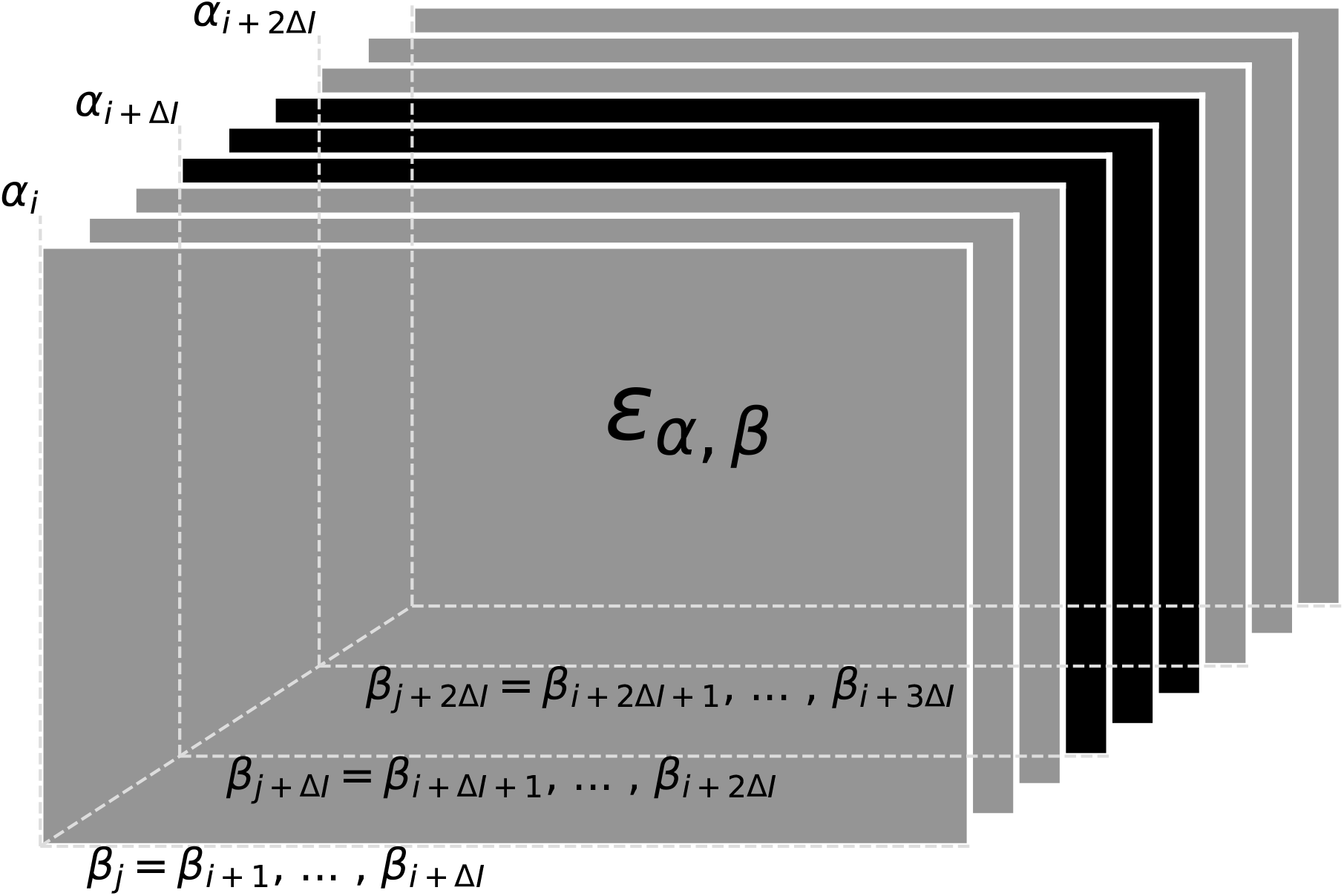
Distance cube containing all distances, *ε_α,β_* Eq. (3), for possible signal pairs *α* and *β* within the current tracking window. Each layer, referring to a time step i, contains the distances between all signals *α_i_* detected at this time and their potential signal partners *β_j_* detected maximally 10 s after signal *α_i_* (Δ*I* time-steps after *i*). Distances in grey layers correspond to signal pairs where one signal partner could potentially have a smaller distance to a signal outside the error cube. Only connections based on the distances in the central black layers can be assumed to be valid, since all potential connections of both signal partners are within the error cube. Connections established for the black layers are assigned to signal traces obtained in previous tracking steps in a second step.

**Figure 5.**
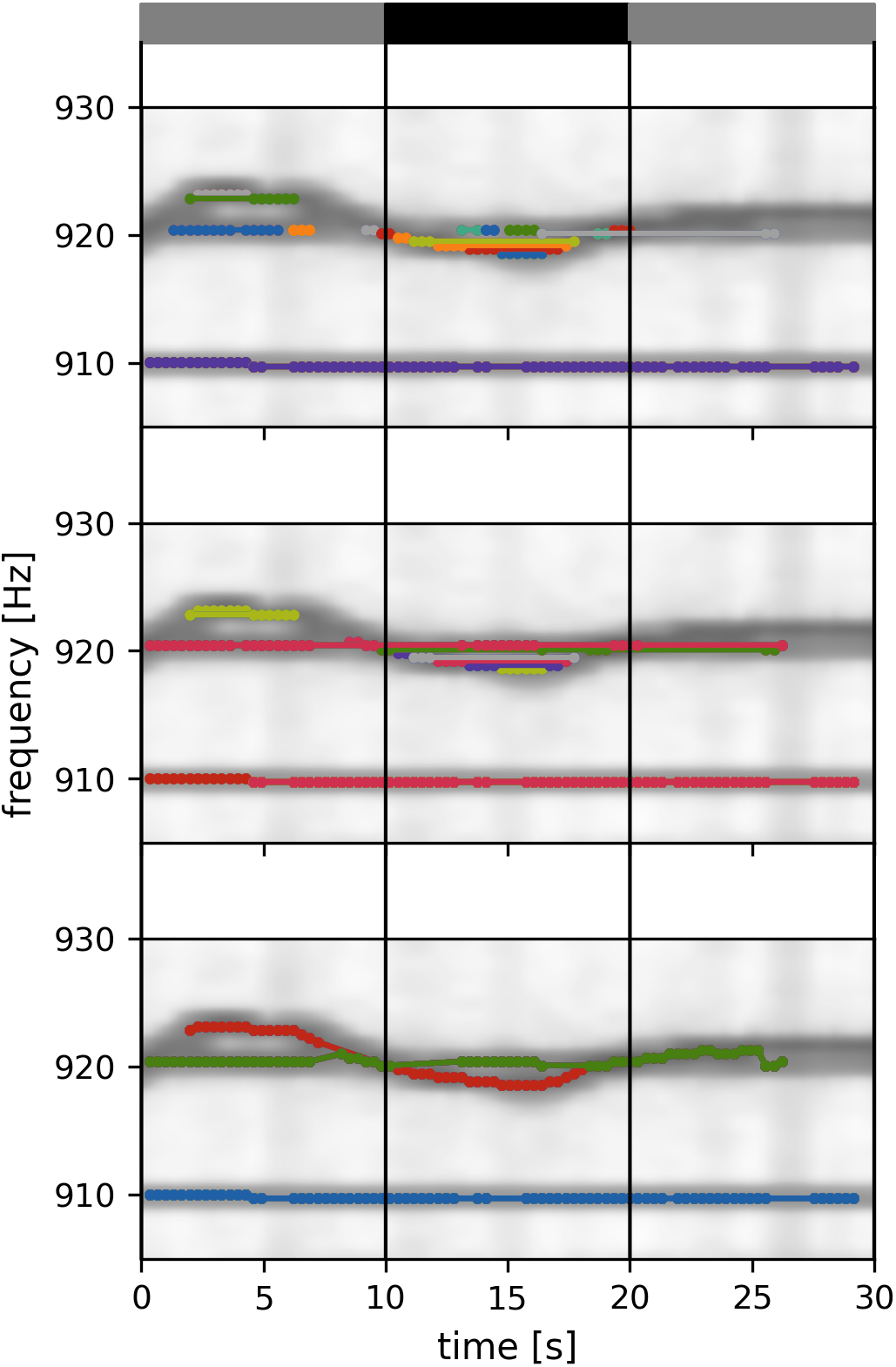
Tracking within a data window. Signals detected in 30 s data window are connected to each other and assigned to fish identities according to their distance *ε*, Eq. (3). Signal pairs with smaller distances are connected first. With increasing distance values, more connections and identities are formed, complemented, or merged, ensuring no temporal overlap. Different stages of this tracking algorithm are shown in the three panels. While multiple separate signal traces are still present in the upper panels (different colors), only three EODf traces corresponding to three fish identities are left upon completion of the algorithm (bottom panel). Only signal pairs within the central 10 s of an 30 s tracking window (vertical lines) are assigned to already established fish identities from previous tracking windows. The summed spectrogram of a 30 s long part of the recording shown in Fig. 2B is shown in the background.

Signal pairs are then connected and assigned to potential fish identities based on the values in the distance cube. The algorithm described in the following (Fig. 5) is a kind of clustering algorithm that has a notion of temporal sequence. The resulting clusters are traces of different fish identities (“labels”) tracked over time.

The signal pairs are traversed in order of ascending distances. If one of *α_i_* or *β_j_* have already been assigned to a fish identity, then this pair is added to this fish identity. If *α_i_* coincides with one fish identity and *β_j_* with another one, then the two fish identities are merged. If neither *α_i_* nor *β_j_* match an existing fish identity, the pair is assigned to a new fish identity. Assignment to or merging of fish identities are only possible in the absence of temporal conflicts, i.e. a fish identity cannot have more than one signal at the same time. In case of temporal conflicts, the signal pair is ignored and the algorithm proceeds with the next one. As a result, we obtain signal traces built upon minimal signal errors within a 30 seconds tracking window (Fig. 5).

Since signals within the first and last 10 seconds of a tracking window could have lower distances to signals outside the current tracking window, these connections are potentially flawed (grey layers in Fig. 4; grey bars in Fig. 5). Only connections established within the central 10 seconds take all other potential signal partners into account. Accordingly, only the section of assembled signal traces corresponding to these central 10 seconds of the current tracking window is considered for further processing, where the signal traces are appended to already validated, previously detected ones (Fig. 6).

**Figure 6.**
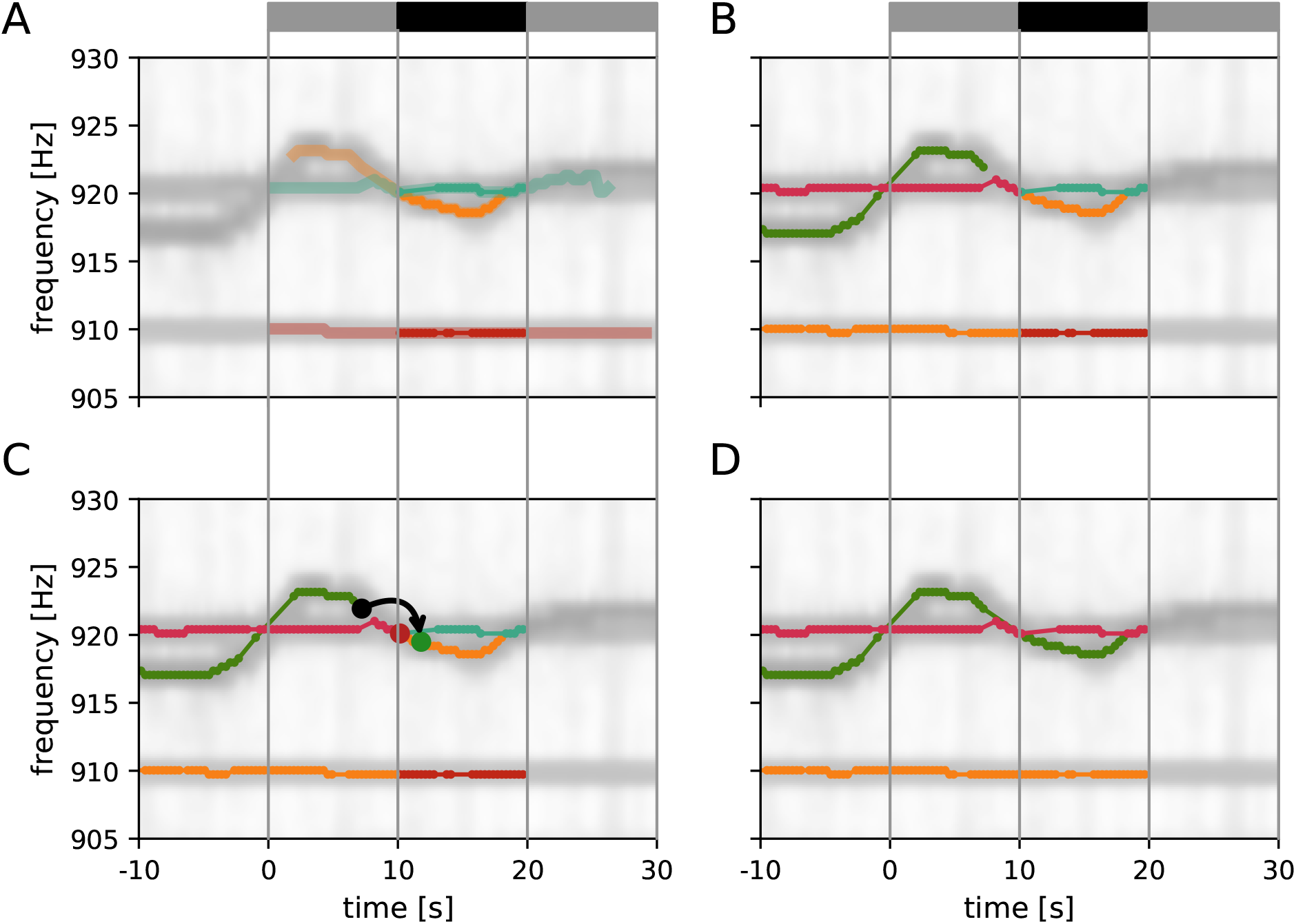
Assembly of tracking results over data windows. **A** New fish identities established within the current tracking window (gray and black bar on top)). Only the central 10 s of these EODf traces (solid traces; black bar) can be assumed to be valid since signals before and after (transparent traces; grey bars) have potential signal partners outside the tracking window. **B** Additional display of EODf traces established in previous iterations if the tracking algorithm. **C** Signal traces are connected according to the smallest possible distance measure between any signal between the last 10 s of the established fish identities (0 < *t* < 10 s) and the central 10 s of the new fish identities (10 s < *t* < 20 s). In the example shown, the distance between the origin signal (black dor) and the target signal (green dor) is the smallest between these two signal traces, accordingly the two signal traces are merged (green and orange lines). An alternative signal (red dot) has a larger distance to the origin signal. **D** Final result of the tracking algorithm that will be used for the next iteration.

#### 3.1.5 Assembly of tracking results over data windows

The assignment of the 10 second long signal traces obtained by the tracking algorithm from 30 s long data windows (Fig. 6 A) to preceding tracking steps (Fig. 6 B) proceeds, similar to the algorithm described above, based on the smallest distances between them.

First, the distance between those signals *α* within the first 10 seconds of the current tracking window of already established fish identities and new signals *β* from the central 10 seconds of the current tracking window are computed. Then, starting with the pair with the smallest distance, the new signal trace containing signal *β* (for example the green dot in Fig. 6C), is connected to the established signal trace (from previous tracking steps) containing signal *α* (for example, the black dot in Fig. 6C). This step is repeated with signal pairs of increasing distance until all possible connections are established (Fig. 6D).

The described tracking within a data window and the subsequent assignment to previously established fish identities is repeated with data windows shifted by 10 seconds until the end of the recording is reached. In each iteration, the distance cube is updated. The first layers corresponding to the first 10 seconds of the previous tracking window are removed (frontal grey layers in fig. 4) and new layers for the next 10 seconds beyond the last tracking window are extended to the error cube in preparation for the next iteration of tracking.

### 3.2 GUI for checking and correcting tracking results

Even though the introduced tracking algorithm is capable of tracking EODs of wave-type electric fish with unprecedented accuracy (see below), occasional tracking errors still remain. We developed a GUI that allows to visually inspect and validate tracked EODf traces and to fix flawed connections (Fig. 7).

**Figure 7.**
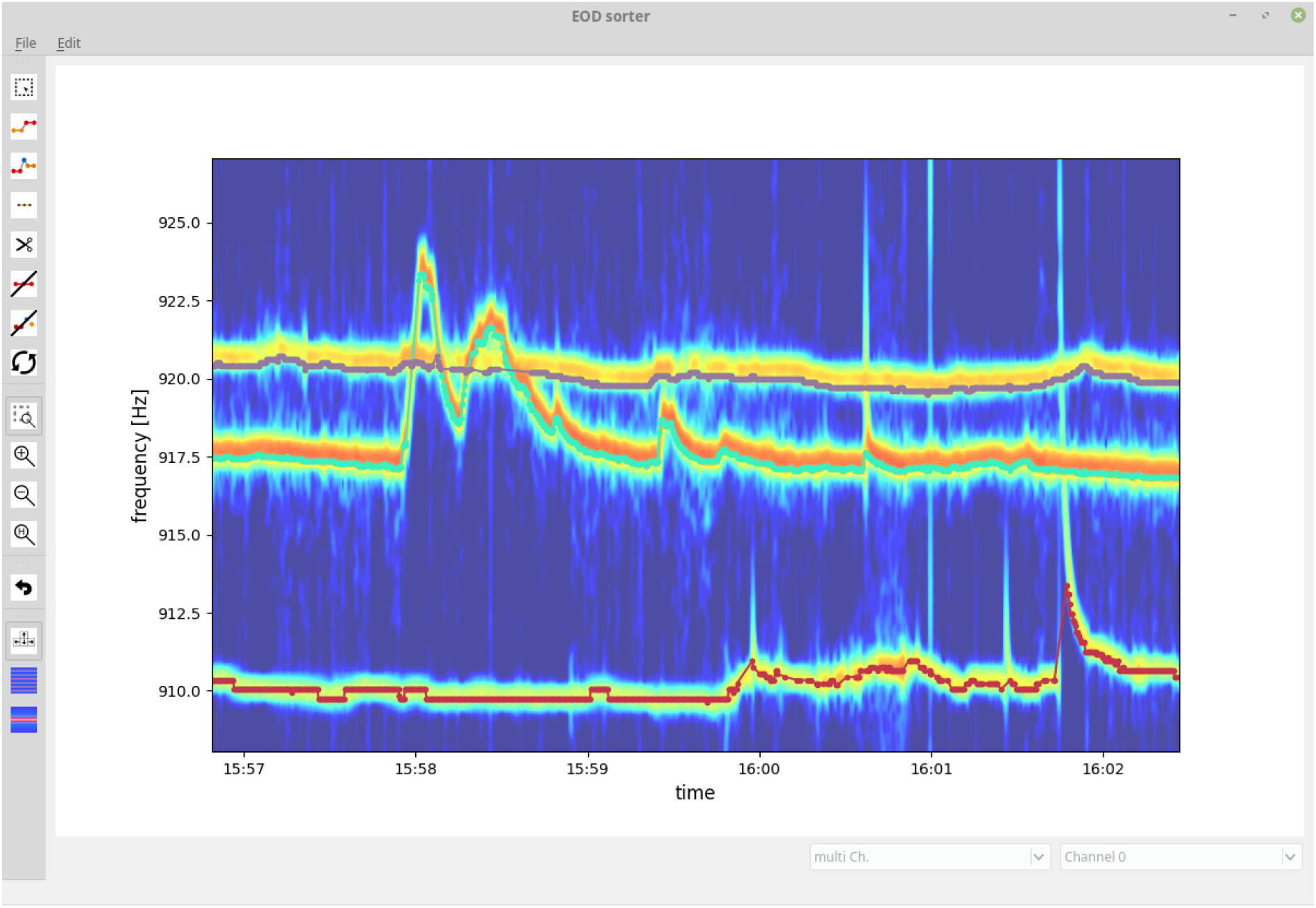
Graphical user interface for validating and fixing tracking results. The user is presented with the tracked signal traces (EODf traces) displayed on top of a spectrogram summed up across recording electrodes. The user can delete, cut, and connect signal traces or delete signals not originating from electric fish.

Flawed connections can easily be identified by their clear deviation from the spectrogram displayed in the background. Furthermore, signal traces with a detection gap beyond the temporal threshold of Δ*t_thresh_ =* 10 s of the tracking algorithm can be manually connected based on visual cues from the spectrogram. The resulting validated signal traces are then stored and further analyzed (Raab et al., 2019, 2021).

### 3.3 Performance of the tracking algorithm

In order to quantify the performance of the presented tracking algorithm, we tracked signals of a whole population of *A. leptorhynchus* recorded with an 8 × 8 electrode array in a small stream in the Llanos of Colombia during the day of April 10*^th^*, 2016 for 10h:50m. First, we tracked the fish with the presented algorithm and then visually inspected, corrected, and validated the tracking results using the GUI (Fig. 7). Second, we run the tracking algorithm again and compared the connections made by the algorithm with the manually improved ones. That is, for each signal *α_i_* we inspected all possible connections with a signal *β_j_* (one row in the distance cube) within the central 10 seconds of the current tracking window. If all the *β_j_* for a given *α_i_* were assigned to the same fish identity in the visually corrected tracking results, we have no potential conflict and these connections were not further considered for quantifying the performance of the algorithm, because these are the simple cases with a single fish within the maximum EODf difference, Δ*f_thresh_*, of 2.5 Hz. If, however, the possible connections involved two or more fish identities, a tracking conflict was possible. For each such potential tracking conflict, we extracted the EODf difference Δ*f*, Eq. (2), field difference Δ*S*, Eq. (6), frequency error *ε_f_*, Eq. (4), field error *ε_S_*, Eq. (7), and resulting distance measure *ε*, Eq. (3), between the signal *α_i_* and the best signal partner *β_j_*, the one with the smallest distance ε, associated with the same fish identity as in the visually corrected signal traces (true connection), as well as between the signal *α_i_* and the best signal partner *β_j_* belonging to a different fish identity (false connection). Further fish identities of the *β_j_* with larger distances were ignored.

In order to assess the performance of each signal feature difference (Δ*f* & Δ*S*) and distance measure (*ε_f_, ε_S_, ε*) in separating true from false connections, we computed the fraction of signal differences or errors of true connections being smaller than those of the corresponding false connections. If this fraction would be 100 % then the tracking algorithm would always have connected the right signals. In addition we quantified the overlap of the two distributions by the area under the curve (AUC) of a receiver-operating characteristic (ROC). Despite an overlap (low AUC values) in principle 100 % correct connections would be possible, but an overlap demonstrates that fixed decision thresholds are not feasible.

We start with evaluating the 464 tracking conflicts from a 5 minute snippet being especially challenging to track, because of several crossings of EODf traces (Fig. 8). A small EODf range of this 5 minute data snippet is displayed in Fig. 7. The least reliable tracking feature appears to be the difference in EODf (Δ*f* and *ε_f_*). Frequency differences of true connections were smaller than the ones of false connections in only 94.83 % (440/464) of the cases (Fig. 8 A, C). Better results can be achieved based on the field error (Δ*S* and *ε_S_*) as a tracking feature. The field differences of true connections were smaller in 99.57 % (462/464) of the cases (Fig. 8 B, D). However, this performance can even be improved when using the distance measure, ε, that combines both the frequency error, *ε_f_*, and field error, *ε_S_*. In 99.87 % (462/464) of the tracking conflicts, true connections had smaller distances than false connections (Fig. 8 E). The AUC values for all measures were similar to the fractions of correct connections (Δ*f* and *ε_f_*: AUC= 95.16 %, Δ*S* and *ε_S_*: AUC= 99.77 %, *ε*: AUC= 99.86 %), indicating a small but existing overlap between the two distributions.

**Figure 8.**
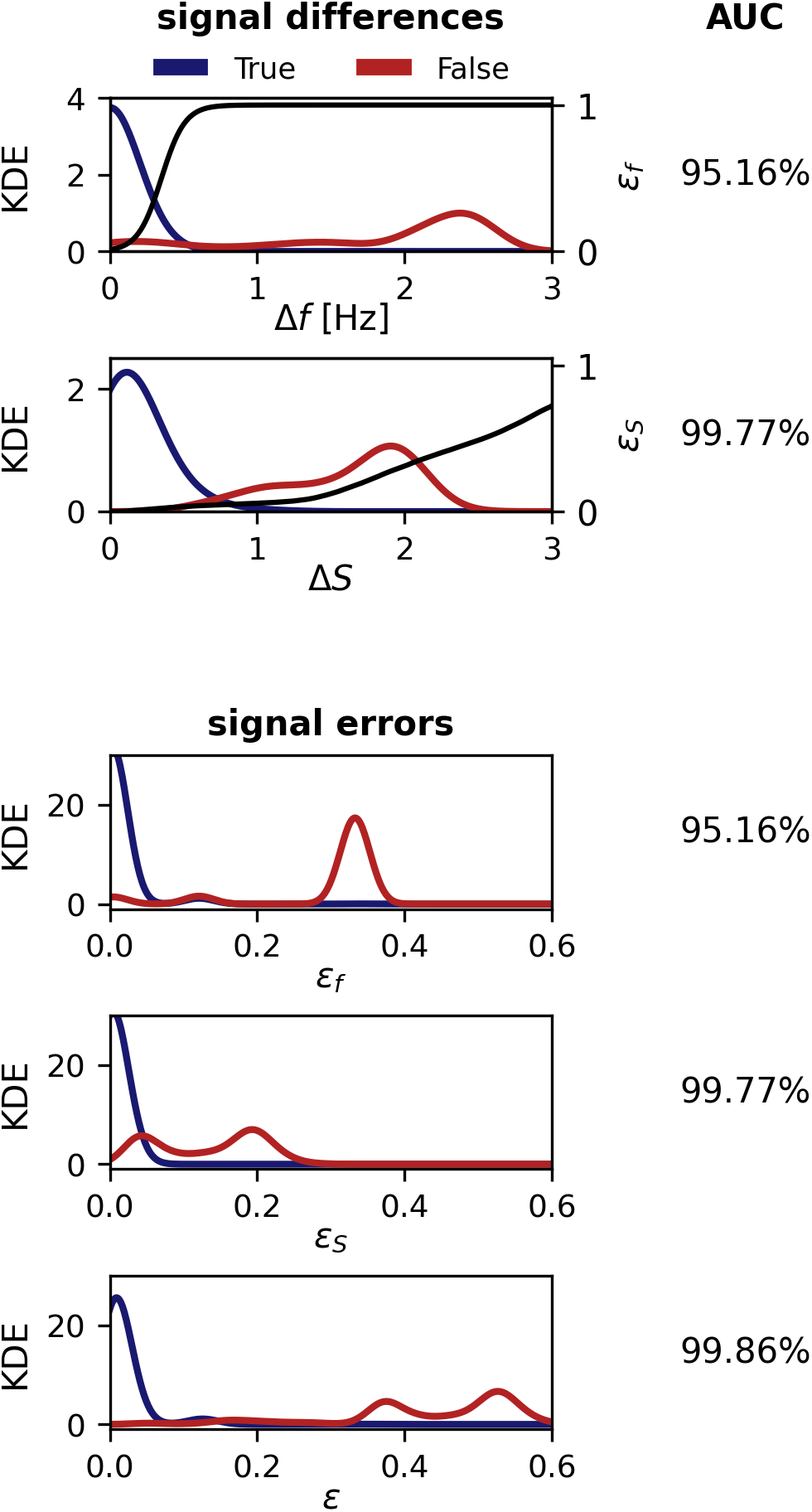
Performance of the tracking algorithm. Conflicts appear if signals could be connected to multiple different fish identities, that have been manually corrected and checked *post hoc* (fig. 7). In most but not all cases, correct connections have smaller signal differences or errors (red) than wrong connections (orange). Shown are kernel density estimates (KDE) for the various signal differences, errors, and distances. The overlap of the distributions was quantified by the AUC of an ROC-analysis as indicated in the right column. **A** EODf differences, Δ*f* Eq. (2). A logistic function, Eq. (4) (black line), translates EODf difference to frequency errors, *ε_f_*. **B** Field differences, Δ*S*, Eq. (6). The cumulative distribution (black line) of field differences of all pairings, not only from conflicts, translates field differences to field errors, *ε_S_*, Eq. (7). **C** Frequency error, *ε_f_*, Eq. (4). **D** field error, **E** Combined distance measure, *ε*, Eq. (3). Note, that frequency and field errors (C, D) are mapped via monotnoically increasing functions from signal differences (A, B) and thus result in the same fraction of correct connections and AUC values. However, the distance measure combining both field and frequency error performs best.

The 261 344 tracking conflicts of the whole recording yield similar results. However, the higher proportion of “easy” tracking conflicts increased the performance of the various features in general and differences between them were less pronounced. Nevertheless, EODf still performed worse (99.73 % correct connections) than field difference (99.81 % correct connections). Again, combining both into the distance measure, Eq. (3), resulted in the best performance (99.95 % correct connections). Correspondingly, the overlap between the two distributions was reduced (Δ*f* and *ε_f_*: AUC= 99.79 %, Δ*S* and *ε_S_*: AUC= 99.85 %, *ε*: AUC= 99.98 %).

## 4 RESULTS

The complexity of the data set we recorded in Colombia in 2016 led us to the development of the presented tracking algorithm. The high density of fish in this data set (about 25 fish within 3.5 × 3.5 m^2^) results in more similar individual EODf traces that frequently cross each other, in particular in the context of communication (fig. 9). This severely challenged previous tracking approaches (Madhav et al., 2018; Henninger et al., 2020), thus a better tracking algorithm was required. The improved algorithm resolves many tracking issues resulting from crossing EODf traces and facilitates the evaluation of wave-type electric fish recordings even in abundant populations.

**Figure 9.**
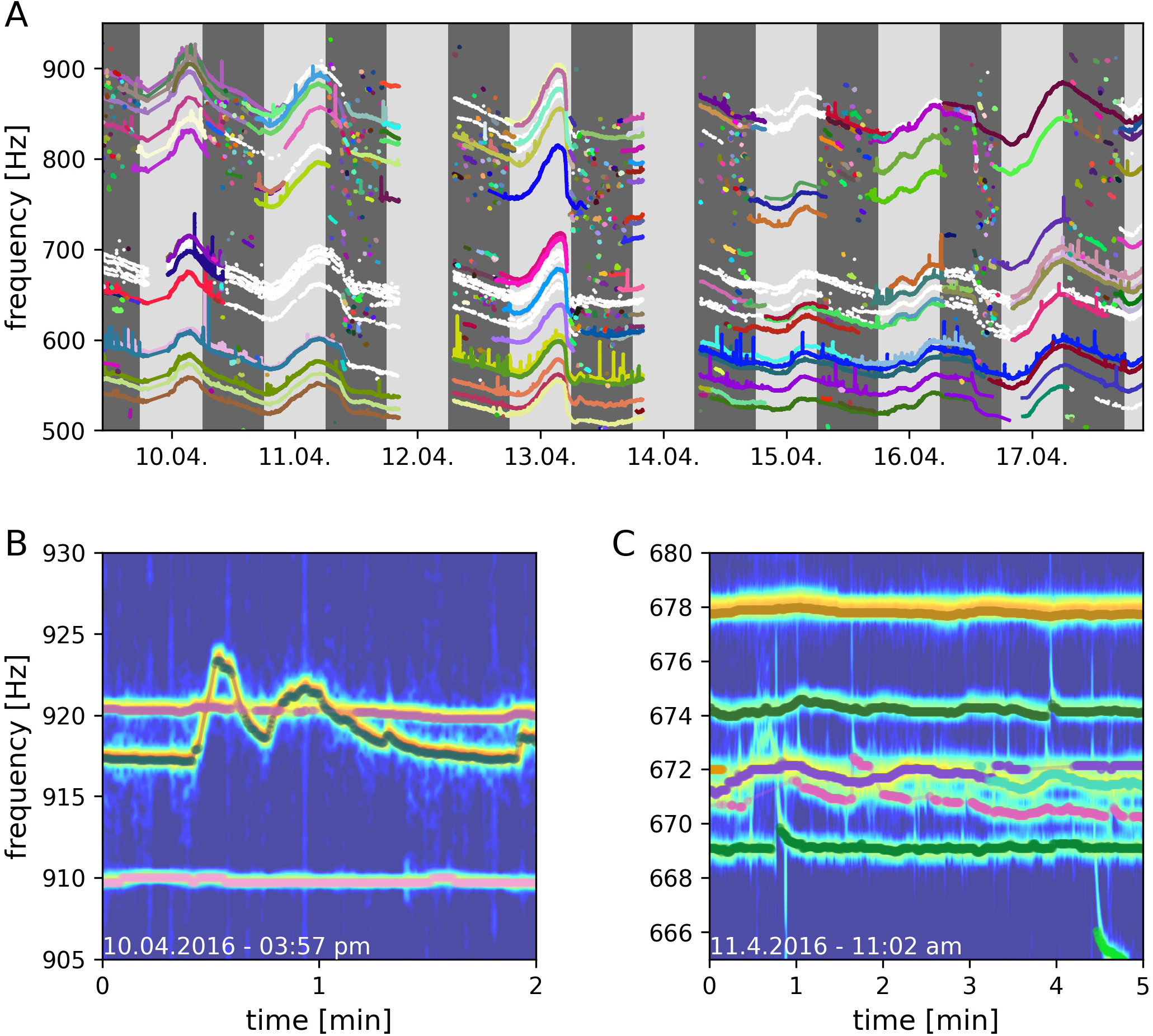
Long-term field recording of A. *macrostomus*, a member of the A. *leptorhynchus* species group, in Colombia, 2016. EODs were recorded with a 64 channel electrode array covering 3.5 × 3.5 m^3^. **A** Eight days of detected and tracked EODfs. Successfully tracked and validated signal traced of different fish are indicated in different colors. Signal traces that could not be clearly validated are indicated in white. Dark gray areas indicate night time, light gray areas day time. **B** Signal traced of three fish where the crossing EODf traces of the upper two fish could reliably be resolved by the tracking algorithm. **C** Too many signal traces with similar frequencies compromise the tracking algorithm (670 – 672 Hz). Frequency peaks in PSDs belonging to multiple fish temporally overlay and prevent successful tracking.

By means of the developed algorithm we were able, for the first time, to track electric signals of individual fish for multiple consecutive days in a natural, high density population of *A. leptorhynchus* recorded in a stream in Colombia (fig. 9). This allowed for novel insights into the natural behavior of these fish in the wild, including their communication and movement behaviors. A preliminary analysis of the tracked fish indicate that many fish stay pretty stationary within distinct areas for multiple days (Fig. 10). Other fish, especially during the night, can only be tracked for short time periods, suggesting these fish to only transit through the area covered by the electrode array (fig. 9A). Furthermore, fish seem to interact with each other by modulating their EOD frequency in various ways and on many different time scales ranging from seconds to many minutes, if not even hours. This includes not only distinct communication signals like rises (Raab et al., 2021, fig. 9B), but also other not yet classified EODf modulations, for example multiple EODf traces entwining each other (fig. 9C). Such natural observations are invaluable since they represent the ground-truth of natural and undisturbed behaviors a given animal species evolved to. Only in the wild, the whole scope of an animal’s behavior can be observed in the context of all relevant stimuli and conditions that shaped these behavior through evolutionary adaptations. Accordingly, such natural observations yield the unique opportunity to discover novel and unexpected behavioral traits and associated causalities. For example, Fortune et al. (2020) described behavioral and physiological adaptations of *Eigenmannia vicentespelea*, another gymnotiform wave-type electric fish, in response to living in a constantly dark cave. *E. vicentespelea* developed increased territoriality and enhanced EOD amplitudes in comparison to *Eigenmannia trilineata*, not living in caves, to face the challenges of their specific habitat. If not for the corresponding field study, these behavioral and physiological adaptations probably never would have been discovered.

**Figure 10.**
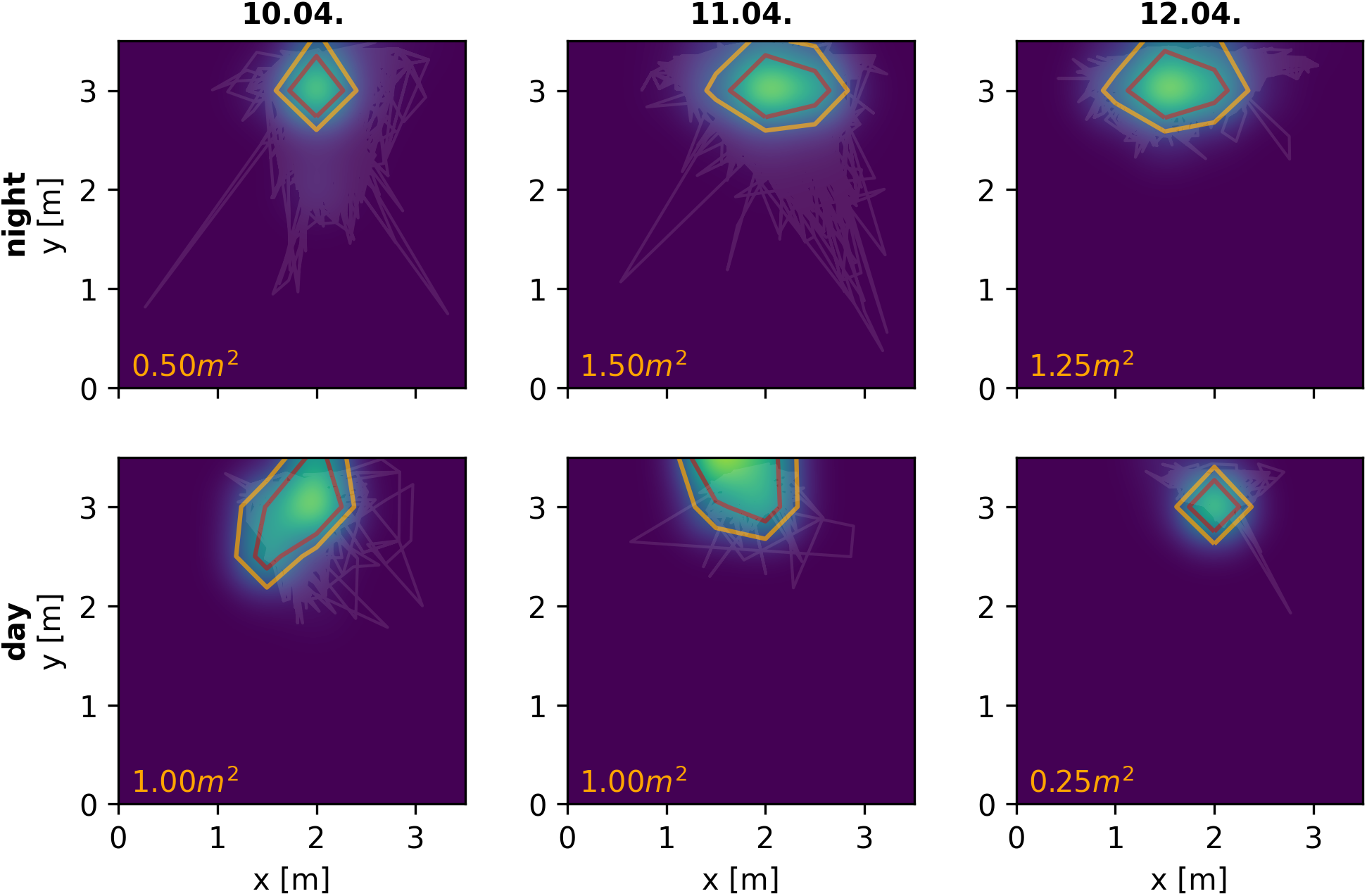
Spatial behavior of a single *A. macrostomus* detected and tracked consecutively for four days. Heat-maps and contour lines show the fish’s probability of presence across the monitored 3.5 × 3.5 m^2^ area of the river during the night (top panels) and day (bottom panels). Orange contour lines include the area in which the fish spends more than 50 % of the time, the red lines more than 75 % of the time respectively. Even though the fish certainly shows movement behaviors, especially during the night, it remains remarkably stationary in the bottom-right corner of the observation area for four consecutive days.

Furthermore, field studies are also essential to validate conclusions drawn form laboratory experiments, which is especially important since behaviors observed in the laboratory often deviate from those observed in the wild (Cheney et al., 1995; Rendall et al., 1999; Henninger et al., 2018). In our case, the preliminary behavioral obsevations we made in Colombia and described above fit to and support the conclusions of our recent laboratory experiments (Raab et al., 2019, 2021). In these experiments we used the here presented algorithm to track individual electric signals of *A. leptorhynchus* in different behavioral contexts. This includes the evaluation of individual spatio-temporal movement behaviors in a freely moving and interacting group of 14 *A. leptorhynchus* (Raab et al., 2019) as well as the communication behavior of pairs of *A. leptorhynchus* competing over a shelter during staged competitions (Raab et al., 2021). In both laboratory and field observations fish produce the majority of rises as electrocommunication signals during the night (Raab et al., 2021, Fig. 9 A, B), are more stationary during the day compared to the night (Raab et al., 2019, Fig. 10), and seem to not remain completely stationary for the whole inactive day-phase but rather show short periods of activity (Raab et al., 2019, Fig. 10). The observed stationarity of fish observed in our field recordings also fit to our suggestion of *A. leptorhynchus* establishing a dominance hierarchy to regulate an individual’s access to resources (Raab et al., 2021). Due to the fish’s stationarity, repetitive conflicts with the same individuals are presumably inevitable and the establishment of a dominance hierarchy can be assumed to be the most economic way to resolve these conflicts (Sapolsky, 2005).

Finally, the evaluation of natural recordings very accurately illustrate the advantages and limitations of the presented algorithm. While crossing EODf traces can usually be resolved accurately (fig. 9B), reliable tracking usually fails when to many signals traces are of similar EODf and entwine in diffuse EODf alterations (fig. 9C). In these occasions the signals of multiple fish superimpose in the spectrogram analysis (Fig. 2C) for longer time periods. Accordingly, after the EODf traces disentangle later on the clear assignment to one of the involved identities is usually impossible.

## 5 DISCUSSION

Previous approaches on tracking wave-type EODs of individual wave-type electric fish either utilized their EODf (Henninger et al., 2020) or the spatial profile of their electric fields (Madhav et al., 2018) as tracking features. We assessed the performance of both signal features alone as well as a combination of both, based on tracking conflicts occurring while processing a recording of a natural, high density population of *A. leptorhynchus* in a stream in Colombia. The comparison of spatial field properties clearly performs better than a comparison of EOD frequencies. Certainly, the EODf of *A. leptorhynchus* can be remarkably stable over minutes to hours (Moortgat et al., 1998). However, EODf can also change on many size and time scales, because of its strong temperature dependence (Dunlap et al., 2000), actively produced electrocommunication signals (e.g. Zupanc, 2002; Triefenbach and Zakon, 2008; Smith, 2013; Benda, 2020; Raab et al., 2021), and also as an artifact of the EODf extraction from the PSDs (fig. 2). Accordingly, the suitability of EODf as tracking feature decreases the more fish are recorded and analyzed simultaneously, since EODf differences between fish are potentially smaller and interactions between fish involving active EODf modulations can be assumed to be more frequent. Therefore, spatial field properties reflecting a fish’s spatial position and orientation represent a more robust tracking feature, especially when only those signal pairs with small EODf differences are considered for comparison and tracking.

The best tracking performance is achieved by using both EODf differences and field differences. This combined signal distance implements a tracking bias that helps to resolve tracking conflicts in at least two scenarios, where tracking solely based on field differences fail. First, if two fish swim close to each other with similar orientations, then their spatial profiles are similar but they can be still differentiated based on their EODfs. Second, in the event of crossing EODf traces. In such intersections, temporarily only one signal can be extracted by detecting peaks in the PSD (Fig. 3A). So neither the EODf difference nor the difference in spatial profiles provide a meaningful hint for tracking in the moment of the intersection. Adding EODf difference to the distance measure then slightly favors connections of signal pairs with more similar EODfs, resulting in a bias for superimposed signals detected at the intersection to be connected to the EODf trace of the fish with a more constant EODf (fig. 5, green trace in bottom panel). The other signal traces, accordingly, remain to be connected across the intersection afterwards.

More important for the improved performance of the presented tracking algorithm is the algorithm itself, in addition to the combined distance measure. The tracking algorithm establishes connections within an extended tracking window based on smallest distances (fig. 5). This is in contrast to existing tracking algorithms (Madhav et al., 2018; Henninger et al., 2018, 2020), that immediately connect the signals detected in a given time step to known fish identities.

When studying animals and their behaviors by means of evaluating external recordings, we rely on the detection of sensory cues emitted actively or passively by the animals themselves (Dell et al., 2014; Hughey et al., 2018). In cluttered environments or when signals are weak (low signal-to-noise-ratio), reliable signal detection is often impaired and detection losses frequently occur. These detection gaps complicate reliable tracking, especially when signals are tracked according to their temporal occurrence. In recordings of electric fish, detection losses frequently result from fish being too far away from recording electrodes, from the electric fields being blocked by any objects between a fish and recording electrodes, or by intersections of EODfs. The resulting tracking failures can be avoided with the presented algorithm, since it relies less on the temporal sequence of detected signals. Rather, connections are established according to the smallest distances within extended tracking windows.

Despite the high fractions of correct connections (fig. 8), the resulting EODf traces need to be corrected manually. This is in particular necessary in sections with close by EODfs or EODf modulations that frequently cross other EODfs. Recordings of only a few fish with well separated EODfs require much less or even no manual interventions. With the current state of the presented algorithm we push the limits to more complicated signal interactions, but the performance still is not perfect for interesting scenes with a lot of interactions (fig. 9). Deep neural networks that are successfully used to track animal pose (e.g. Mathis et al., 2018), or to annotate acoustic signals from various animals (e.g. Steinfath et al., 2021), might be an interesting option to further improve tracking performance. Such approaches, however, require extensive training data sets. Our tracking algorithm and evaluated data sets might set the basis for developing and training of deep neural networks in the future.

## 6 CONCLUSION

The self-generated electric fields of electric fish offer an unique opportunity for studying natural movement and communication behaviors of nocturnal fish in freely interacting populations. The EODs of whole groups can be recorded simultaneously by means of electrode arrays submerged in the water — without the need to catch and tag the fish. The presented algorithm for tracking wave-type electric fish combines previous approaches based on either the individual-specific EODf (Henninger et al., 2020) or the spatial profiles of electric fields resulting from a fish’s location and orientation (Madhav et al., 2018). The algorithm uses a compound signal distance, that incorporates both EODf and spatial profiles. We developed a new temporal clustering method that assembles fish identities from all signals within a large tracking window according to ascending signal distances. With this approach, our algorithm improves in resolving tracking issues mainly resulting from crossing EODf traces or detection losses, and enables tracking of wave-type EODs with unprecedented accuracy. Since tracked EOD traces allow insights into both movement and communication behaviors, a reliable tracking algorithm is key to many behavioral studies, both in the laboratory and in the field, that have not been possible before. From such big-data behavioral studies we expect many novel insights into the sensory ecology and into social and communication behaviors of these fascinating fishes (Henninger et al., 2018; Fortune et al., 2020; Raab et al., 2019, 2021), that also impact the way we study sensory processing.

## CONTRIBUTIONS

The algorithm described and presented in this manuscript was developed by TR but builds upon collective or individual ideas shared between all Coauthors. MM, RJP, and NJC developed the idea to utilize spatial electric field properties of individual electric fish as a feature to track their EODs (Madhav et al., 2018). JH and JB provided the tools required for detecting and extracting electric signals of wave-type electric fish in multi-electrode grid recordings and developed the first algorithmic approaches to track EODs of electric fish by means of their individual specific EOD frequency (Henninger et al., 2020). TR developed the here presented tracking algorithm which combines and refines previous approaches. This includes the implementation of the combined signal distance, which considers both electric field difference and EOD frequency, to determine signal similarities, as well as the idea to tack signals according to their similarity (combined signal distance) in discrete tracking windows. TR, furthermore, developed software tools to inspect and post-process tracked EOD traces, illustrated and evaluated the algorithm, and wrote the manuscript. Many of the ideas realized in the presented algorithm originated from collaborations and numerous discussion of TR with the other Authors.

